# Comparative Reproductive Biology And Impacts Of Selfing In Two Epiphytic Bromeliaceae

**DOI:** 10.64898/2026.07.20.739642

**Authors:** Joshua M. Felton, M. Shane Heschel, Erin N. Bodine, Rachel S. Jabaily

## Abstract

The breeding systems of most Bromeliaceae have not been tested directly. We characterized the floral biology and breeding systems of two epiphytic, tank-forming bromeliads endemic to the Atlantic Forest of Brazil, *Vriesea rafaelii* (Tillandsioideae) and *Billbergia brasiliensis* (Bromelioideae), using controlled self- and cross-pollinations in a greenhouse. Both species were self-compatible. Based on seed-set, the self-compatibility index was 0.59 in *V. rafaelii* and 0.87 in *B. brasiliensis*. In *B. brasiliensis*, selfed flowers produced lighter fruits and lighter seeds than outcrossed flowers, whereas in *V. rafaelii* fruit and seed mass did not differ between self- and cross-pollinated flowers. A subset of *V. rafaelii* flowers showed a spatial separation of the anthers and stigma (herkogamy) that prevented autonomous selfing. These results add two species to the small set of bromeliads with experimentally characterized breeding systems and highlight differences in reproductive output with selfing.

## Introduction

A plant’s breeding system, whether its flowers are self-compatible or self-incompatible, is a fundamental part of its life history, determining seed set and pollination ecology. Many plants are self compatible and can self-pollinate (Igic & Busch 2013), which can be advantageous when mates or pollinators are scarce because it provides reproductive assurance. However, selfing can be costly due to reductions in heterozygosity, which can result in the expression of deleterious recessive alleles. This often produces inbreeding depression, in which selfed offspring set fewer or lighter fruits and fewer or less viable seeds than outcrossed offspring (Hedrick & Kalinowski 2000; Angeloni et al. 2011). Determining whether a species can self, and whether selfing reduces its reproductive output, contextualizes the evolutionary trends of breeding systems and builds natural history knowledge. The Neotropical family Bromeliaceae is large and varied in floral form, and its breeding systems span the full range from obligate selfing to obligate outcrossing, either through self-incompatibility (SI) mechanisms or through dioecy (reviewed in Cascante-Marín & Núñez-Hidalgo 2023). Self-compatibility is common, occurring in nearly all genera of subfamily Tillandsioideae and in only a few genera of Bromelioideae, the two subfamilies that are the focus of this study. Some species also vary in flower size and in the arrangement of their reproductive organs within a single inflorescence, with certain flowers favoring outcrossing through a spatial separation of the anthers and stigma (herkogamy). For most bromeliads, however, the breeding system has never been directly tested (Cascante-Marín & Núñez-Hidalgo 2023). Here we characterize and compare the reproductive biology of two epiphytic, tank-forming bromeliads endemic to the Atlantic Forest of Brazil. *Vriesea rafaelii* Leme (subfamily Tillandsioideae) produces an erect spike inflorescence and dry, capsular fruits, while *Billbergia brasiliensis* L.B.Sm. (subfamily Bromelioideae) produces a pendant spike inflorescence and fleshy fruits. We (a) describe the reproductive biology of each species, including variation among individual flowers, and (b) test whether the biomass a plant invests in fruits and seeds differs between selfed and outcrossed flowers.

## Methods

The taxa were purchased from Tropiflora (Sarasota, FL, USA) and grown from seed collected from a single parental plant of each species. Plant care and earlier developmental data are described in (Jabaily et al. 2021). Seventeen *V. rafaelii* and 14 *B. brasiliensis* individuals with visible developing inflorescences were included in the study. Within each species, individuals were assigned to a selfing or an outcrossing treatment (*V. rafaelii*: 9 self, 8 outcross; *B. brasiliensis*: 7 self, 7 outcross). Flowers left unmanipulated were scored as autonomously selfed. *V. rafaelii* is protogynous: the stigma becomes receptive shortly after the petals open, and the anthers dehisce about three hours later. We monitored receptivity with a hand lens, noting receptivity once the stigma was visibly wet with secretions from its surface papillae. For the outcrossing treatment, pollen was applied to receptive stigmas in the evening, and the anthers were removed before dehiscence to prevent self-pollination. When only one flower was available on an individual (N = 9 flowers), the excised anthers were stored at −20°C and used later as donor pollen, as frozen bromeliad pollen remains viable for crossing (De Souza et al. 2015). *Billbergia brasiliensis* anthers and stigma mature together and remain in proximity during anthesis. Pollinations were done in late morning to coincide with anther dehiscence. For the outcrossing treatment, flowers were opened by hand before the petals curled back, the anthers were removed to prevent accidental self-pollination, and the flower was enclosed in a fine mesh organza bag, which helped ensure no accidental crossing occurred from pollen from other plants, until the controlled cross was performed.

Fruits of *B. brasiliensis* and *V. rafaelii* require roughly seven and eighteen months, respectively, to reach full maturity (a color change from green to yellow in *B. brasiliensis*, and the onset of capsule dehiscence in *V. rafaelii*). Because of time constraints on data collection, inflorescences were harvested before full ripeness, after about four months for *B. brasiliensis* and six months for *V. rafaelii*. We considered a flower aborted if it failed to produce any seed (seed mass = 0 g). Harvested inflorescences were dried in paper bags for five days at 60°C, and the rachis, scape bracts, and floral bracts were weighed separately from the fruits. Each dried fruit was dissected to separate seeds and their associated structures from fruit tissue, and seeds and fruit were each weighed on an analytical balance, with seed mass recorded as the total mass of all seeds per fruit rather than the mass of an individual seed. To quantify each species’ capacity for self-fertilization, we calculated a Self-Compatibility Index (Zapata & Arroyo 1978).

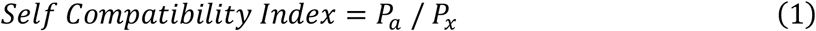

where *P*_*a*_ is the proportion of selfed flowers that set fruit and *P*_*x*_ is the proportion of cross-pollinated flowers that set fruit. The index ranges from 0 (completely self-incompatible) to 1 (fully self-compatible). The index assumes Pa ≤ Px; values above 1 would reflect experimental error rather than genuine self-compatibility. To test whether cross-type (selfed vs. outcrossed) influenced fruit mass or seed mass, we used a nested mixed-model ANOVA with species, cross-type, and their interaction as fixed effects and individual plant identity nested within species as a random effect. Analyses were performed in JMP (SAS Institute).

## Results

### Natural history

*Billbergia brasiliensis* had a spike inflorescence that was pendant and white from farinose hairs (Figure 1A). Stigma receptivity occurred at the same time of anther dehiscence which had occurred by the time the corolla has spiraled into a coil (Figure 1B-C). Flowers did not exhibit any form of herkogamy (Figure 1C). The typical showy pink bracts varied in pigment between individuals. Multiple individuals had bracts that were brown and dried at anthesis (Figure 1D-E). Multiple individual scapes did not elongate enough to become pendant and instead flowered when erect, primarily within the tubular vegetative rosette. Flowers on these scapes were difficult to observe and manipulate. Fruits changed pigment from green to yellow once ripe (at least six months post anthesis) and were subrotund with ribs. Seeds were coated in a mucilaginous substance (Figure 1F).

**FIGURE 1.**
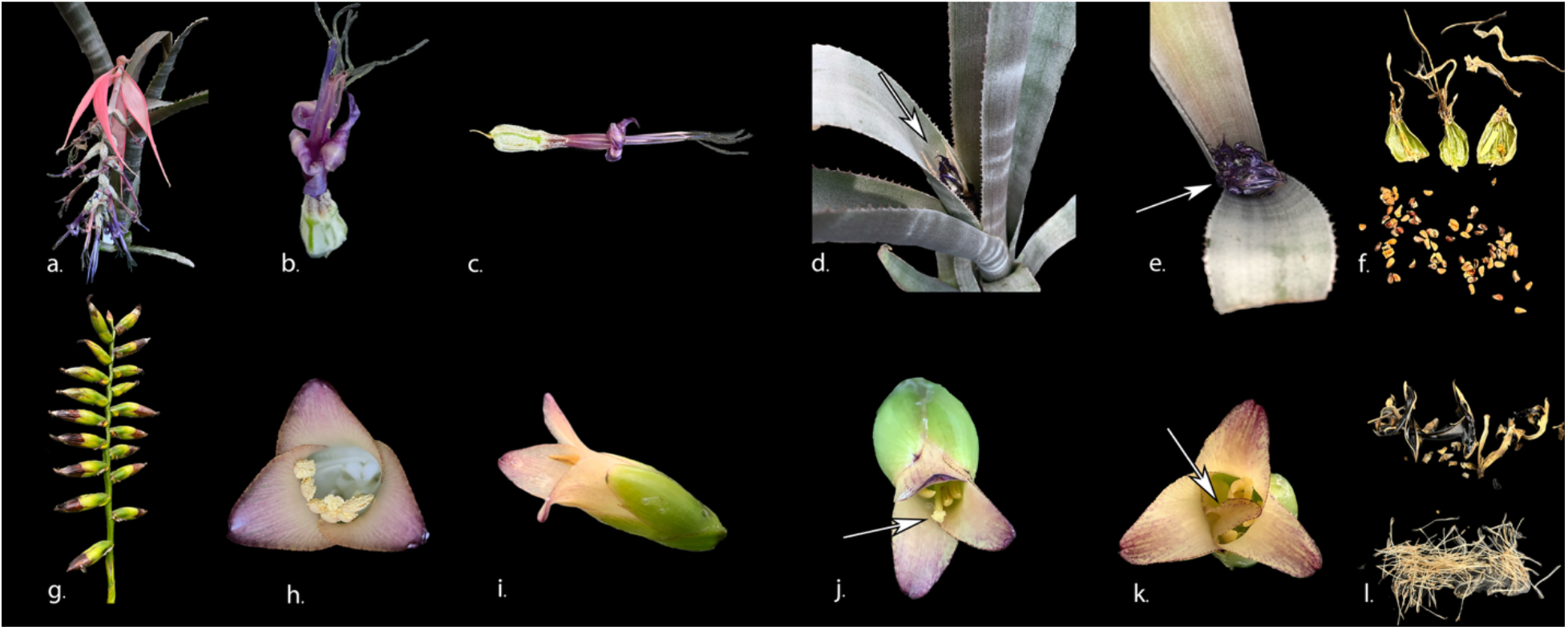
Reproductive morphology of *Billbergia brasiliensis* (a–f) and *Vriesea rafaelii* (g–l). **A**. Pendant spike inflorescence with showy pink floral bracts, the axis white with farinose hairs. **B**. Single flower at anthesis (ca. 65 mm long; Smith & Downs 1979) with the corolla coiled, the stigma receptive as the anthers dehisce. **C**. Open flower (sepals ca. 10 mm long; Smith & Downs 1979) showing the anthers and stigma in contact, with no spatial separation of the two (no herkogamy); stamens are shorter than the petals in this species, consistent with the observed contact. **D. E**. Individuals whose scapes did not elongate, so the inflorescence flowered erect within the leaf rosette, with floral bracts brown and dried at anthesis (arrows). **F**. Ripe fruits, which turn from green to yellow and are subrotund and ribbed, and seeds coated in mucilage. **G**. Erect spike inflorescence with dark green scape and floral bracts. **H**. Flower in face view (petals ca. 48 × 30 mm; Leme 1999), the anthers held against the stigma. **I**. Flower in lateral view (flower ca. 65 mm long; Leme 1999); anthers dehisce about three hours after the petals open. **J**. Flower in which the stamens are shorter than the pistil (arrow), so self-pollination did not occur. **K**. Flower in which a petal appendage (cymbiform, ca. 12–14 × 6 mm; Leme 1999) extends across the stigma (arrow), preventing self-pollination. **L**. Dehisced capsules and plumose seeds at maturity, about seventeen months after flowering.

*Vriesea rafaelii* had an erect spike inflorescence with dark green scape and floral bracts (Figure 1G). Stigma receptivity began shortly after petals had opened, with anthers dehiscing approximately three hours after petal opening (Figure 1I). Anthers were arranged ventrally in the flower, touching the stigma (Figure 1H). Most individuals had anthers at the same length as the stigma. Mismatch in anther and pistil length was noted particularly in individuals (*N* = 10 individuals) who began their development later in the study population. The most common mismatch observed was the stamen being shorter than the pistil length, so much so that selfing did not occur (*N* = 29 flowers) (Figure 1J). Petal appendages, which are a single flap of tissue between the staminal filament and the petal at the base of the corolla (Brown & Terry 1992), occasionally were long enough to cover the stigma (*N* = 4 flowers), leading to no fruit set from selfing (Figure 1K). Capsule development took about seventeen months from flowering (Figure 1L).

### Breeding system

All hand-pollinated flowers formed fruits regardless of cross type. However, successful seed production was lower following selfing than outcrossing (81% vs. 93% in *B. brasiliensis* and 34% vs. 58% in *V. rafaelii*), indicating that the primary reproductive cost of selfing occurred after fruit initiation (Table 1). The effect of cross type on capsule mass and on seed mass did not differ significantly between the two species (capsule mass, F = 1.29, df = 1, P = 0.257; seed mass, F = 1.19, df = 1, P = 0.276), so we report each species separately. In *Billbergia brasiliensis*, outcrossed flowers produced heavier fruits (F = 4.08, df = 1, P = 0.045) and heavier seeds (F = 7.23, df = 1, P = 0.008) than selfed flowers (Figure 2A). In *Vriesea rafaelii*, capsule mass was marginally lower in outcrossed than selfed flowers but did not differ significantly between cross types (F = 2.48, df = 1, P = 0.116; Table S1), and seed mass did not differ between cross types when flowers that set no seed are included (F = 0.85, df = 1, P = 0.356; Table S2; Figure 2B). Both species were self-compatible, with a self-compatibility index of 0.87 in *B. brasiliensis* and 0.59 in *V. rafaelii* (Table 2).

**FIGURE 2.**
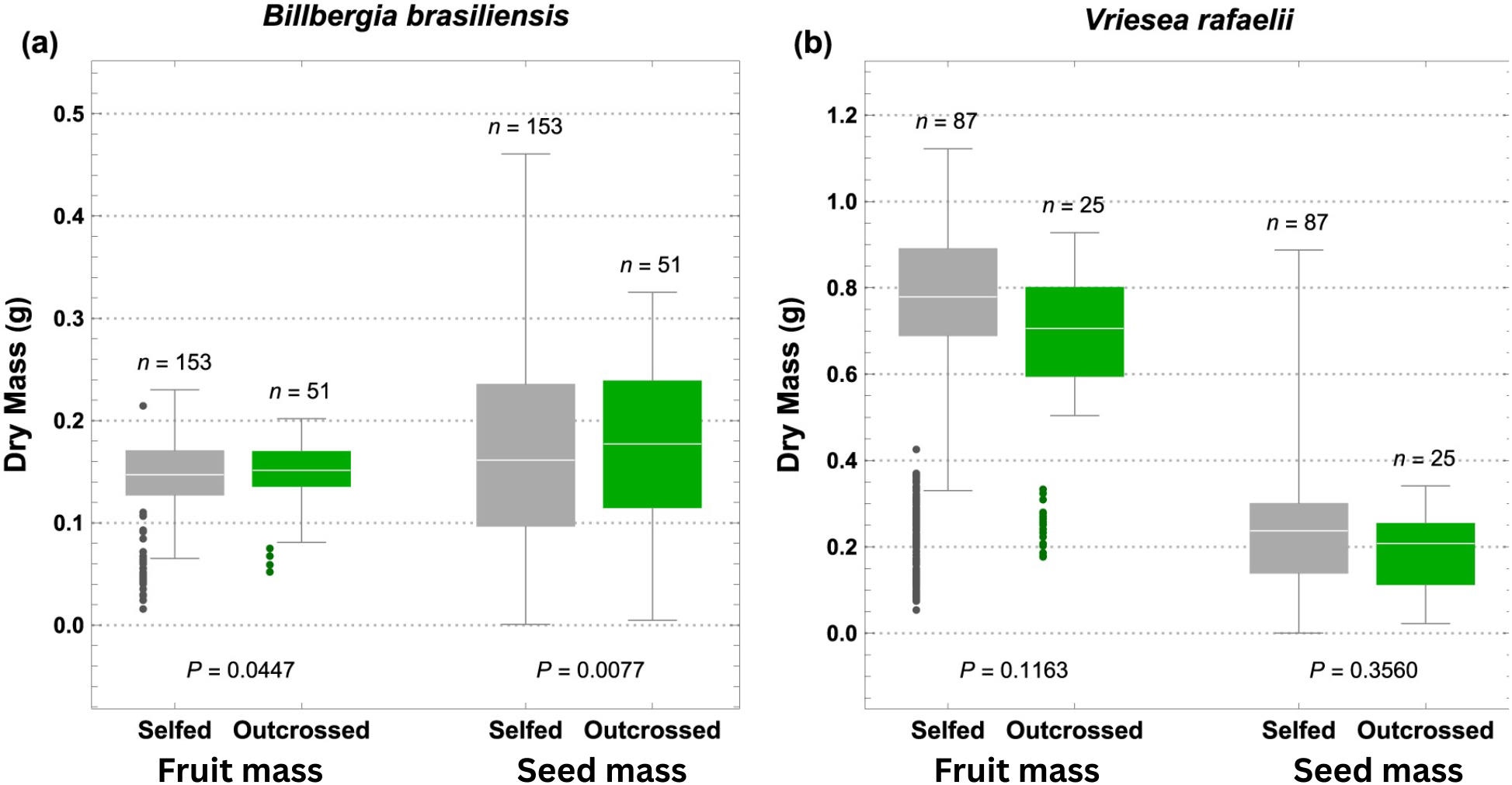
Dry mass of fruits and seeds from selfed and outcrossed flowers of **A**. *Billbergia brasiliensis* and **B**. *Vriesea rafaelii*. In each panel the left pair of boxes is fruit mass, and the right pair is total seed mass including appendages; selfed flowers are shown in grey and outcrossed flowers in green. The number of flowers (n) is given above each box. P values are for the effect of cross type within each species, from the mixed-model ANOVAs in Table S1 (fruit mass) and Table S2 (seed mass). The points show the fruit mass of aborted flowers (*B. brasiliensis*: 37 selfed, 4 outcrossed; *V. rafaelii*: 166 selfed, 18 outcrossed).

**TABLE 1.**
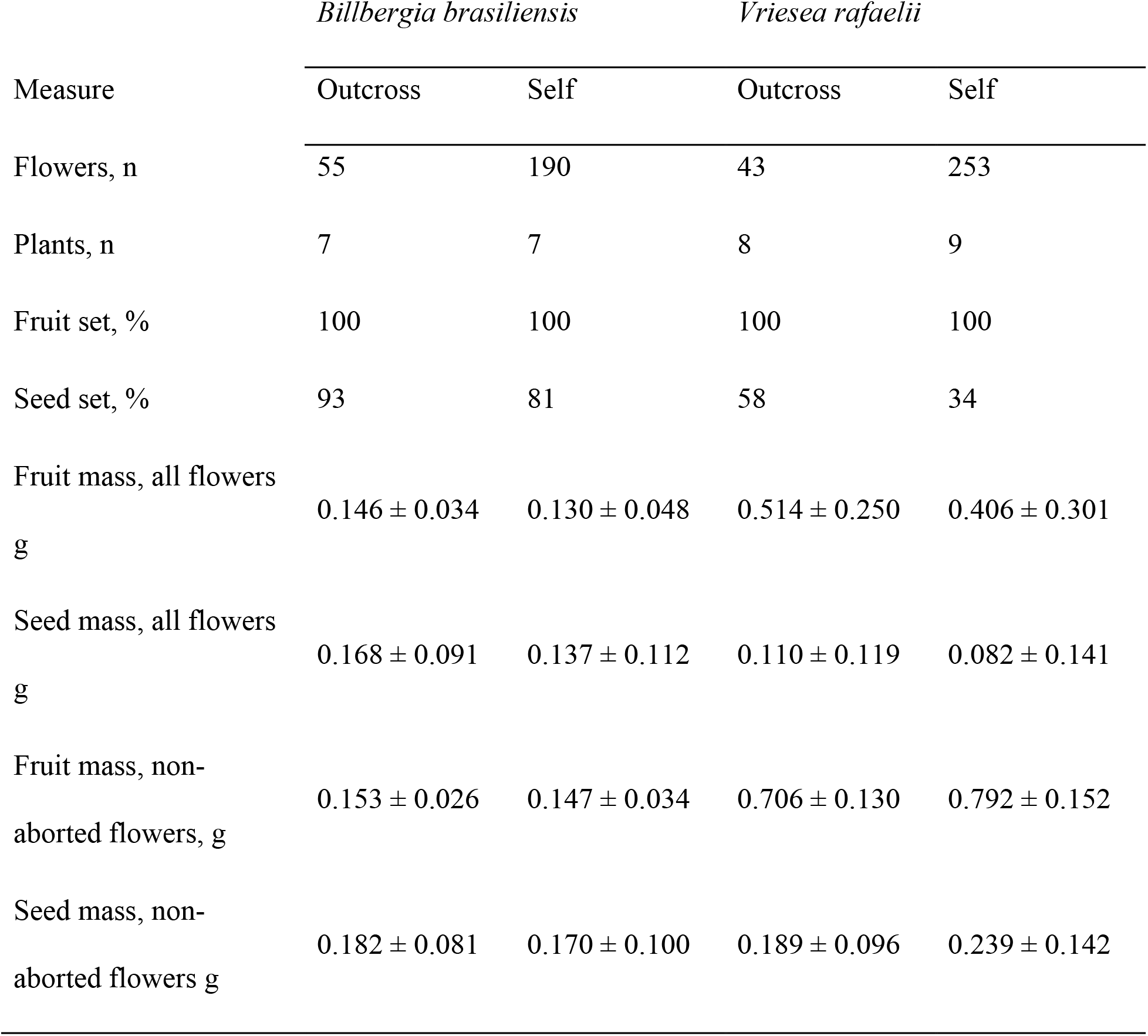
Breeding-system descriptors for hand-pollinated flowers of *Billbergia brasiliensis* and *Vriesea rafaelii*. Fruit and seed mass are mean ± SD. Fruit set is the proportion of flowers forming a capsule; seed set is the proportion producing seed. A flower was scored as aborted if it failed to set any seed (seed mass = 0 g). “All flowers” rows include aborted flowers; “non-aborted flowers” rows exclude them and correspond to the values plotted in Fig. 2 and analyzed in Tables S1–S2.

| Measure | *Billbergia brasiliensis* |  | *Vriesea rafaellii* |  |
| --- | --- | --- | --- | --- |
|  | Outcross | Self | Outcross | Self |
| Flowers, n | 55 | 190 | 43 | 253 |
| Plants, n | 7 | 7 | 8 | 9 |
| Fruit set, % | 100 | 100 | 100 | 100 |
| Seed set, % | 93 | 81 | 58 | 34 |
| Fruit mass, all flowers<br>g | 0.146 ± 0.034 | 0.130 ± 0.048 | 0.514 ± 0.250 | 0.406 ± 0.301 |
| Seed mass, all flowers<br>g | 0.168 ± 0.091 | 0.137 ± 0.112 | 0.110 ± 0.119 | 0.082 ± 0.141 |
| Fruit mass, non-aborted flowers, g | 0.153 ± 0.026 | 0.147 ± 0.034 | 0.706 ± 0.130 | 0.792 ± 0.152 |
| Seed mass, non-aborted flowers g | 0.182 ± 0.081 | 0.170 ± 0.100 | 0.189 ± 0.096 | 0.239 ± 0.142 |

**TABLE 2.** Indices of self-compatibility and inbreeding depression for *B. brasiliensis* and *V. rafaelii*. SC-index (Eq. 1) calculated as selfed set ÷ outcrossed set; values near 1 indicate self-compatibility and near 0 self-incompatibility, with the conventional threshold for self-compatibility at approximately 0.2 (Zapata & Arroyo 1978).

| Index | *Billbergia brasiliensis* | *Vriesea raphaelii* |
| --- | --- | --- |
| SC-index (fruit set) | 1.00 | 1.00 |
| SC-index (seed set) | 0.87 | 0.59 |

## Discussion

Both species were self-compatible, but selfing was costlier in more ways in *B. brasiliensis* than in *V. rafaelii*. In *B. brasiliensis*, selfed flowers produced lighter fruits and lighter seeds than outcrossed flowers and set seed somewhat less often, indicating early-acting inbreeding depression in both the formation and the provisioning of seed. Reduced fruit set and smaller seeds following self-pollination have likewise been reported in field studies of *Canistrum aurantiacum* and *Encholirium heloisae* (Christianini et al. 2013)(Siqueira Filho & Machado 2001), placing *B. brasiliensis* among the bromeliads in which selfing lowers early reproductive output. In *V. rafaelii*, the cost of selfing was confined to a single step. Fruit and seed mass did not differ between cross types, but selfed flowers were far less likely to set any seed at all than outcrossed flowers (34% versus 58%). Among the flowers that did set seed, the seed crop was no smaller after selfing than after outcrossing. The lower reproductive output of selfed *V. rafaelii* therefore came largely from flowers that failed to set seed, which might be indicative of self-compatibility issues and higher abortion rates due to inbreeding. Mean seed mass alone therefore understated inbreeding depression in *V. rafaelii*, because much of the effect was carried by the many selfed flowers that set no seed. A subset of *V. rafaelii* flowers was also herkogamous, with the anthers held away from the stigma or a petal appendage covering it, and these flowers formed fruits and set seed only when outcrossed, although this was observed in only three flowers. Because fruit and seed mass index an early stage of offspring performance, these measures bound rather than resolve the total strength of inbreeding depression in either species; resolving it will require following selfed and outcrossed progeny to germination and later life-history stages (Riginos et al. 2007) as well as estimating inbreeding directly with molecular markers (Matos Paggi et al. 2015). These results add two previously uncharacterized species to the small set of bromeliads with experimentally tested breeding systems (Cascante-Marín & Núñez-Hidalgo 2023). Because both species can self yet incur a cost at the seed stage, selfing and inbreeding may impact population dynamics. However, the degree to which selfing contributes to population persistence and recruitment in the wild remains an open question that greenhouse crosses can only begin to answer.

Both species fell well within the self-compatible range by an index of self-incompatibility, though *V. rafaelii* was the less self-compatible of the two (Table 2). Our finding runs counter to the general expectation for the family: self-compatibility is thought to be widespread in Tillandsioideae and comparatively rare in Bromelioideae (Cascante-Marín & Núñez-Hidalgo 2023), yet here the bromelioid *B. brasiliensis* was more self-compatible than the tillandsioid *V. rafaelii*. This suggests that breeding system in Bromeliaceae can vary even within lineages generally considered self-incompatible, and underscores how little direct testing exists outside a handful of species. The within-plant variation in herkogamy in *V. rafaelii* also shows that the capacity to self can vary among flowers on a single individual, which is relevant to how selfing rates are estimated from whole-plant or whole-population data (Whitehead et al. 2018). Self-compatibility promotes reproductive assurance in the absence of mates or pollinators, but selfing carried a measurable cost in both species, expressed as reduced seed set in *V. rafaelii* and as both reduced seed set and lighter seeds in *B. brasiliensis*. Therefore, the capacity to self does not translate directly into equivalent reproductive output, indicating inbreeding depression and potential impacts on population dynamics. Future work should investigate this further on three fronts: following selfed and outcrossed progeny beyond seed to germination and establishment, where later-acting inbreeding depression may yet emerge; estimating realized outcrossing rates in natural populations with molecular markers; and documenting the floral visitors and pollination behavior that determine how often outcrossing occurs at these sites. For two narrowly distributed endemics, resolving how much of their reproduction is selfed, and at what cost, would do much to clarify how secure their mating systems leave them.

## Acknowledgements

The authors would like to thank Ali Keller for plant care assistance, Dr. Brad Oberle and Dr. Brian Sidoti for their insights and comments during the project, and the Colorado College Keller family fund and the Hevey fund for financial support.

## Authors Contribution Statement

JMF, MSH, RSJ conceptualized the project; JMF collected the data; JMF, MSH, ENB analyzed the data; JMF, RSJ wrote the original draft; all authors reviewed and edited the final draft

## Notes

### Competing Interest Statement

The authors have declared no competing interest.

## References

Angeloni, F., Ouborg, N. J., and Leimu, R. 2011. Meta-analysis on the association of population size and life history with inbreeding depression in plants. Biological Conservation 144: 35–43. 10.1016/j.biocon.2010.08.016

Brown, G. K. and Terry, R. G. 1992. Petal Appendages in Bromeliaceae. American Journal of Botany 79: 1051–1071. 10.2307/2444915

Cascante-Marín, A. and Núñez-Hidalgo, S. 2023. A Review of Breeding Systems in the Pineapple Family (Bromeliaceae, Poales). The Botanical Review 89: 308–329. 10.1007/s12229-023-09290-0

Christianini, A. V., Forzza, R. C., and Buzato, S. 2013. Divergence on floral traits and vertebrate pollinators of two endemic Encholirium bromeliads. Plant Biology 15: 360–368. 10.1111/j.1438-8677.2012.00649.x

De Souza, E. H., Souza, F. V. D., Rossi, M. L., Brancalleão, N., Da Silva Ledo, C. A., and Martinelli, A. P. 2015. Viability, storage and ultrastructure analysis of Aechmea bicolor (Bromeliaceae) pollen grains, an endemic species to the Atlantic forest. Euphytica 204: 13–28. 10.1007/s10681-014-1273-3

Hedrick, P. W. and Kalinowski, S. T. 2000. Inbreeding Depression in Conservation Biology. Annual Review of Ecology and Systematics 31: 139–162. 10.1146/annurev.ecolsys.31.1.139

Igic, B. and Busch, J. W. 2013. Is self-fertilization an evolutionary dead end?New Phytologist 198: 386–397. 10.1111/nph.12182

Jabaily, R. S., Oberle, B., Fetterly, E. W., Heschel, M. S., Sidoti, B. J., and Bodine, E. N. 2021. Refining Iteroparity with Comparative Morphometric Data in Bromeliaceae. International Journal of Plant Sciences 182: 577–590. 10.1086/715484

Matos Paggi, G., Palma-Silva, C., Bodanese-Zanettini, M. H., Lexer, C., and Bered, F. 2015. Limited pollen flow and high selfing rates toward geographic range limit in an Atlantic forest bromeliad. Flora - Morphology, Distribution, Functional Ecology of Plants 211: 1–10. 10.1016/j.flora.2015.01.001

Riginos, C., Heschel, M. S., and Schmitt, J. 2007. Maternal effects of drought stress and inbreeding in Impatiens capensis (Balsaminaceae). American Journal of Botany 94: 1984–1991. 10.3732/ajb.94.12.1984

Siqueira Filho, J. A. D. and Machado, I. C. S. 2001. Biologia reprodutiva de Canistrum aurantiacum E. Morren (Bromeliaceae) em remanescente da Floresta Atlântica, Nordeste do Brasil. Acta Botanica Brasilica 15: 427–443. 10.1590/S0102-33062001000300011

Whitehead, M. R., Lanfear, R., Mitchell, R. J., and Karron, J. D. 2018. Plant Mating Systems Often Vary Widely Among Populations. Frontiers in Ecology and Evolution 6: 38. 10.3389/fevo.2018.00038

Zapata, T. R. and Arroyo, M. T. K. 1978. Plant Reproductive Ecology of a Secondary Deciduous Tropical Forest in Venezuela. Biotropica 10: 221. 10.2307/2387907

