## Supplemental Tables for "Comparative Reproductive Biology And Impacts Of Selfing In Two Epiphytic Bromeliaceae"

**Table S1.** Mixed-model analysis of variance for capsule (fruit) mass in *Billbergia brasiliensis* and *Vriesea rafaelii*, with cross type (self vs. outcross) as a fixed effect and plant identity as a random effect.

| Species | Source | *df* | SS | MS | *F* | *P* |
| --- | --- | --- | --- | --- | --- | --- |
| *Billbergia brasiliensis* | Cross type | 1 | 0.007 | 0.007 | 4.08 | 0.045 |
|  | Plant identity (random) | 13 | 0.099 | 0.008 | 4.28 | <0.001 |
|  | Residual | 230 | 0.408 | 0.002 | — | — |
| *Vriesea rafaelii* | Cross type | 1 | 0.102 | 0.102 | 2.48 | 0.116 |
|  | Plant identity (random) | 17 | 12.792 | 0.752 | 18.37 | <0.001 |
|  | Residual | 274 | 11.225 | 0.041 | — | — |

Each effect was tested over the residual mean square within species. The model explained 21% of the variance in capsule mass for *B. brasiliensis* (*n* = 245 flowers) and 54% for *V. rafaelii* (*n* = 293 flowers); plant identity accounted for 16% and 52% of the total variance, respectively. Selfed flowers produced significantly lighter capsules in *B. brasiliensis* but not in *V. rafaelii*. Mean capsule mass by cross type is given in Table 1.

**Table S2.** Mixed-model analysis of variance for seed mass in *Billbergia brasiliensis* and *Vriesea rafaelii* (including flowers that set no seed), with cross type (self vs. outcross) as a fixed effect and plant identity as a random effect.

| Species | Source | *df* | SS | MS | *F* | *P* |
| --- | --- | --- | --- | --- | --- | --- |
| *Billbergia brasiliensis* | Cross type | 1 | 0.076 | 0.076 | 7.23 | 0.008 |
|  | Plant identity (random) | 13 | 0.391 | 0.030 | 2.85 | <0.001 |
|  | Residual | 230 | 2.426 | 0.011 | — | — |
| *Vriesea rafaelii* | Cross type | 1 | 0.010 | 0.010 | 0.85 | 0.356 |
|  | Plant identity (random) | 17 | 2.154 | 0.127 | 11.15 | <0.001 |
|  | Residual | 274 | 3.115 | 0.011 | — | — |

Each effect was tested over the residual mean square within species. The model explained 15% of the variance in seed mass for *B. brasiliensis* (*n* = 245 flowers) and 41% for *V. rafaelii* (*n* = 293 flowers); plant identity accounted for 10% and 39% of the total variance, respectively. Selfed flowers produced significantly lighter seeds in *B. brasiliensis*; seed mass did not differ between cross types in *V. rafaelii*. Mean seed mass by cross type is given in Table 1.
